# Identification of unique mutations in SARS-CoV-2 strains isolated from India suggests its attenuated pathotype

**DOI:** 10.1101/2020.06.06.137604

**Authors:** Shubham Gaurav, Shambhavi Pandey, Apurvasinh Puvar, Tejas Shah, Madhvi Joshi, Chaitanya Joshi, Sachin Kumar

**Author notes:** Corresponding authors (Sachin Kumar), (Chaitanya Joshi). Contributed equally to the work (Shubham Gaurav, Shambhavi Pandey, Apurvasinh Puvar, Tejas Shah).

## Abstract

Severe Acute Respiratory Syndrome Coronavirus-2 (SARS-CoV-2), which was first reported in Wuhan, China in November 2019 has developed into a pandemic since March 2020, causing substantial human casualties and economic losses. Studies on SARS-CoV-2 are being carried out at an unprecedented rate to tackle this threat. Genomics studies, in particular, are indispensable to elucidate the dynamic nature of the RNA genome of SARS-CoV-2. RNA viruses are marked by their unique ability to undergo high rates of mutation in their genome, much more frequently than their hosts, which diversifies their strengths qualifying them to elude host immune response and amplify drug resistance. In this study, we sequenced and analyzed the genomic information of the SARS-CoV-2 isolates from two infected Indian patients and explored the possible implications of point mutations in its biology. In addition to multiple point mutations, we found a remarkable similarity between relatively common mutations of 36-nucleotide deletion in ORF8 of SARS-CoV-2. Our results corroborate with the earlier reported 29-nucleotide deletion in SARS, which was frequent during the early stage of human-to-human transmission. The results will be useful to understand the biology of SARS-CoV-2 and itsattenuation for vaccine development.

## Brief report

Severe Acute Respiratory Syndrome Coronavirus-2 (SARS-CoV-2) is the cause of the novel human Corona Virus Disease COVID-19, first reported on November 17^th^, 2019 in Wuhan, China [12]. It has spread rapidly creating a global public health emergency situation in the last few months and the World Health Organization (WHO) on March 11^th^, 2020 declared it a pandemic. Till date, SARS-CoV-2 has infected close to five million people and caused more than 328,000 deaths globally. Although started late, India showed ramping of more than 112,000 reports of SARS-CoV-2 infection including more than 3,400 deaths, as of May 20^th^, 2020. As compared to the other recent coronavirus epidemics SARS, 2003 and MERS (Middle East Respiratory Syndrome), 2012 having mortality rates of 11% [4] and 35% [8], respectively, SARS-CoV-2 has a current mortality rate of about 3.4% [7]. However, the transmissibility of SARS-CoV-2 recorded far greater than the previous coronavirus epidemics.

SARS-CoV-2 has a positive-sense 29.9 kbp RNA genome, encoding 12 open reading frames (ORFs) [6]. It codes for four structural proteins, namely spike glycoprotein (S), envelope glycoprotein (E), membrane glycoprotein (M), and nucleocapsid protein (N) [12]. S protein is a large membrane protein which helps in endocytosis of the viral particle into the host cell by binding to the angiotensin-converting enzyme 2 (ACE2) receptor [15]. E protein is a small viral transmembrane protein which helps in the assembly of the virions by forming ion channels inside the host cell [5]. M protein is the most abundant protein present on the viral membrane, important for morphogenesis and viral assembly [2]. N protein is responsible for the packaging of the viral genome and plays an important role in viral assembly [13]. The genome also codes for 2 overlapping polyproteins, namely polyprotein 1a (PP1a) and polyprotein 1ab (PP1ab); PP1a being a truncated version of PP1ab. These code for 16 non-structural proteins (NSPs) including proteases (3C-like protease and Papain-like protease) and RNA-processing enzymes like RNA-dependent RNA polymerase, Helicase, 3’-5’ Exonuclease, Endoribonuclease, Guanine N7-methyltransferase, 2’O-ribose methyltransferase and ADP ribose phosphatase [12]. In addition to structural and NSPs, SARS-CoV-2 genome also codes for at least two other viroporin candidates (other than the E protein), namely ORF3a and ORF8 [3]. The function and the possible contribution of other ORFs in the viral infection are largely unknown.

There is a global race to sequence the SARS-CoV-2 to understand its variability among the population and identify the unique mutation. Gujarat Biotechnology Research Centre (GBRC), Gujarat India, actively involved in the complete genome sequencing of SAR-CoV-2 strains from India. More than 144 complete genome sequences of SARS-CoV-2 strains from the state have been completed. The samples were collected from the COVID-19 testing center following the guidelines of Indian council of Medical research. The complete genome sequencing of the samples were carried out by the gene specific primer sets using the Ion S5™ next-generation sequencing system (Thermo Fisher Scientific, USA) following manufacturer’s protocol. In this study, we are reporting the unique mutations in the SARS-CoV-2 genome isolates from India.

The two samples from a male and female aged 66 (husband and wife) has been procured from the COVID-19 testing facility in Gujarat, India. Husband and wife contracted infection from their son who got it from local/community transmission. Both were recovered without having severe symptoms of COVID-19.

The complete genome sequence analysis of SARS-CoV-2 isolated from both the patients showed a size of 29902 bp each (accession numbers MT435081 and MT435082), in contrast with 29903 bp size of the Wuhan strain. However, the single nucleotide deletions in the genome of virus isolates from these two patients were at different positions. Furthermore, the sequence revealed a total of ten mutations each across the genome as compared to SARS-CoV-2 strain from Wuhan. The details of the mutations in the Indian SARS-CoV-2 isolates areprovided in Table 1.

**Table 1.** The position of mutation site with respect to the complete genome of SARS-CoV-2. The other column includes, protein pertaining to the mutation site (protein annotation), reference nucleotide at the same position in Wuhan SARS-CoV-2 sequence, the mutated nucleotide (allele nucleotide), and the corresponding amino acid change due to the point mutation (amino acid mutation) in the full length genome sequence of the Indian virus isolates.

| Position | Protein Annotation | Reference Nucleotide | Sample 1 ♂ 66<br>(Acc. No. <u>MT435081</u>) |  | Sample 2 ♀ 66<br>(Acc. No. <u>MT435082</u>) |  |
| --- | --- | --- | --- | --- | --- | --- |
|  |  |  | Allele Nucleotide | Amino Acid Mutation | Allele Nucleotide | Amino Acid Mutation |
| 241 | 5' UTR: 5' UTR | C | T | - | T | - |
| 1059 | ORF1ab | C | T | Thr > Ile | T | Thr > Ile |
| 3037 | ORF1ab-nsp1 | C | T | - | T | - |
| 14408 | ORF1ab-nsp12 | C | T | Pro > Leu | T | Pro > Leu |
| 15371 | ORF1ab-nsp12 | C | T | Thr > Met | C | - |
| 17747 | ORF1ab-nsp13 | C | C | - | T | Pro > Leu |
| 23403 | S | A | G | Asp > Gly | G | Asp > Gly |
| 25563 | ORF3a | G | T | Gln > His | T | Gln > His |
| 28221 | ORF8 | G | T | Glu > * | T | Glu > * |
| 28254 | ORF8 | A | A | - | Del | Ile > FS |
| 28371 | Gene: N | G | T | Ser > Ile | T | Ser > Ile |
| 28549 | Gene: N | A | Del | Arg > FS | A | - |

The genome sequence of SARS-CoV-2 from the virus isolates of two patients in Gujrat, India, were analyzed and compared with the isolate from Wuhan, China. The single nucleotide change in the viral genome could change the encoding amino acid, which might result in the conformational change in the viral structure protein [11]. The genome of virus isolates from both patients showed a point mutation of C241T corresponding to its genome length and lies in the 5’ untranslated region (5’ UTR). It is unlikely that this mutation has any substantial effect on viral replication. The nucleotide change of C1059T in the protein ORF1ab has directed threonine, a polar uncharged amino acid, to mutate into a hydrophobic amino acid, isoleucine. Threonine residues are well known to form hydrogen-bond interactions with surrounding polar residues in the interior of protein structure. The amino acid residues around this threonine at position 265 corresponding to the polyproteinORF1ab correspond to ^262^K-F-D-**T-**F-N-G-E-C-P^271^ suggesting that threonine might form hydrogen bonds with surrounding polar residues. Perhaps it might also assist in the formation of protein bend caused due to glycine and proline having the position of i, i+3 with respect to each other. Isoleucine being hydrophobic might disturb any conformational protein bend due to its inability to form a hydrogen bond. This mutation might also result in increased hydrophobic interactions with surrounding phenylalanine residues. Since very little is known about NSP2 or any of its homologous structure, no assertion at the structural level could be made.

In the native form, NSP12 is bound with the cofactors NSP7 and NSP8 to form the RNA dependent RNA polymerase (RdRP) complex. The mutation of C14408T at NSP12 in Indian isolates of SARS-CoV-2 lead to the substitution of proline with threonine. The proline residue at position 323 corresponding to the polyprotein results in an unusual turn in the alpha-helix in which it resides [14]. Interestingly, the α-helix is one of the sites for the interaction of NSP12 with the cofactor, NSP8. However, this mutation of proline causes significant changes into the structure due to the disturbance of that turn in that α-helix (Fig. 1A). This would probably change the entire structure of that interacting site, leading to the decreased affinity of RdRP to NSP8 which likely disturbs its functioning. Mutation study in the 3-D structure of NSP12 (Fig. 1B and C) suggests that this mutation results in substantial stabilization of this region (ΔΔG = 1.540 kcal/mol). This mutation also imparts great rigidity to the site (ΔΔS_Vib_ = - 4.074 kcal.mol^-1^.K^-1^).However, the large amount of flexibility originally present in this site might be vital for the functionality of RdRP. This corroborates with our prediction that this mutation might be deleterious for the activity of RdRP.

**Figure 1.**
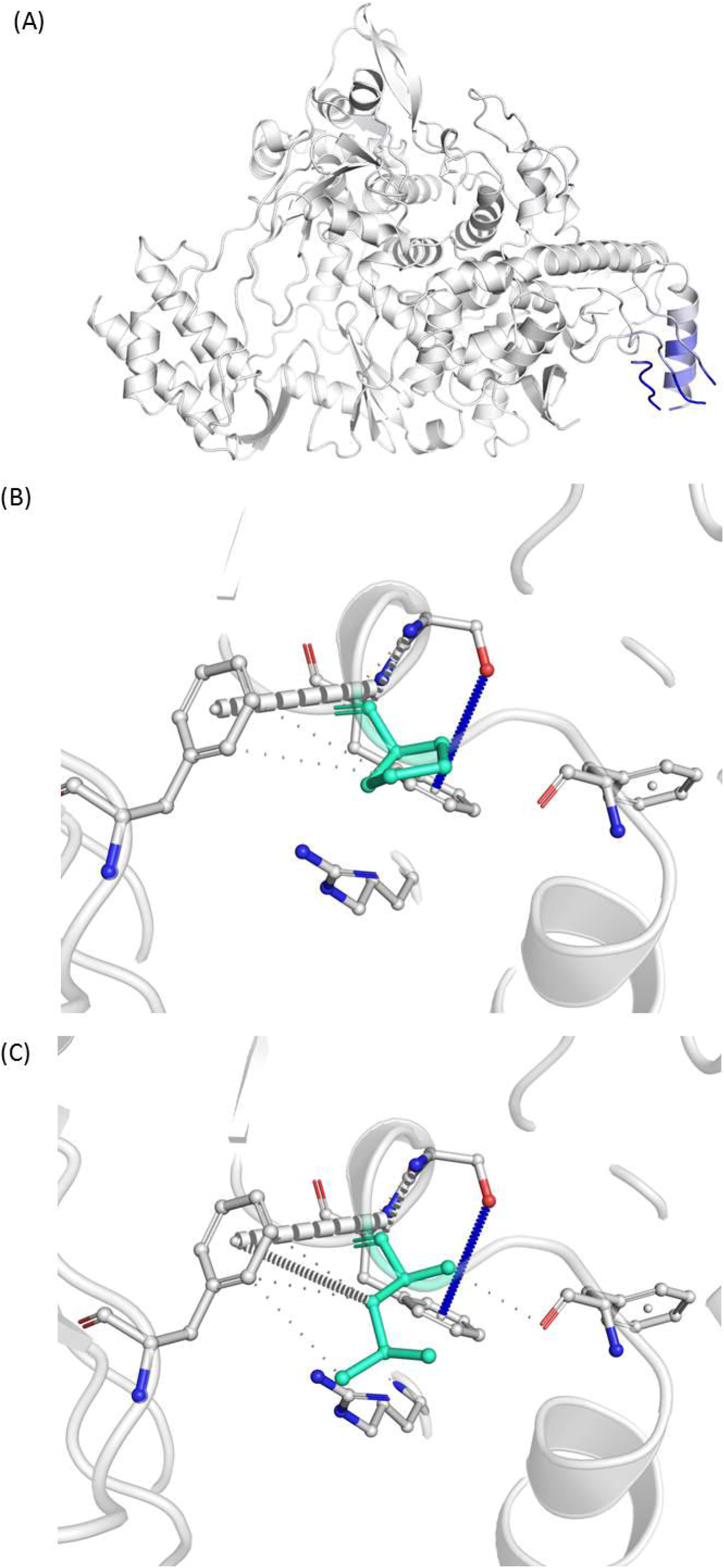
(a) The substitution of proline with leucine at 323^rd^ residue of RdRP substantially changes the inter-atomic interaction of the amino acid residues around the mutation site. The effect of mutation on structure of RdRP has been shown where blue represents a rigidity in the structure and red represents a gain in flexibility. (b and c) The inter-atomic interactions of the residues in vicinity of 323^rd^ amino acid of wild-type RdRP protein and P323L mutated RdRP protein, respectively.

The mutation of polar residue threonine into a hydrophobic residue methionine due to the mutation of C15371T might also significantly affect the protein structure of NSP12. The governing residues around the mutated threonine at position 644 of polyprotein NSP12 are ^640^R-K-H-T-**T**-C-C-S-L-S^649^. Presence of two cysteine residues at an immediate position to threonine propounds formation of cystine residue by the aid of cellular oxidation. The catalytic and enzymatic activity of cysteine residues bound by disulfide bridges plays a crucial role in the folding and stability of extracellular proteins exposed to the harsh extracellular environment [10]. Mutation of threonine into methionine is likely to disturb this disulfide bridge and enzymatic function of cysteine residues, hence the three-dimensional structure and stability of the protein owing to the bulky side chain of methionine [1]. Methionine being a non-polar hydrophobic residue would bury inside the protein thus hampering the immediate cysteine residues in their reactivity as well. Since this amino acid residue lies within a flexible loop region at a site far from the catalytic site of RdRP, this mutation is unlikely to have any significant change in the structureof RdRP.

Mutation of C17747T of viral mRNA results in the substitution of proline into leucine at the 504^th^ amino acid of NSP13 (helicase). However, this site lies in a flexible loop region, far from the functionally active domains of the helicase protein. This mutation is less likely to have any effect on the activity of helicase.

Mutation of A23403G causes the substitution of aspartic acid with glycine. This mutation in the S protein lies in the small loop turn between two anti-parallel beta-strands. These beta-strands participate actively in the trimerization of spike protein which is vital for binding of S protein to its receptor ACE2 [14]. Replacing the aspartic acid residue with the much more flexible glycine might end up in disturbing the β-strands. Prediction of effect of the mutation on the secondary structure (Fig. 2 A) suggests that this mutation is likely to disrupt one of the four beta-strands in that region, thus possibly inhibiting the trimerization of spike protein on the surface of the virus. This could significantly decrease the affinity of S protein to ACE2 receptors as well.

**Figure 2.**
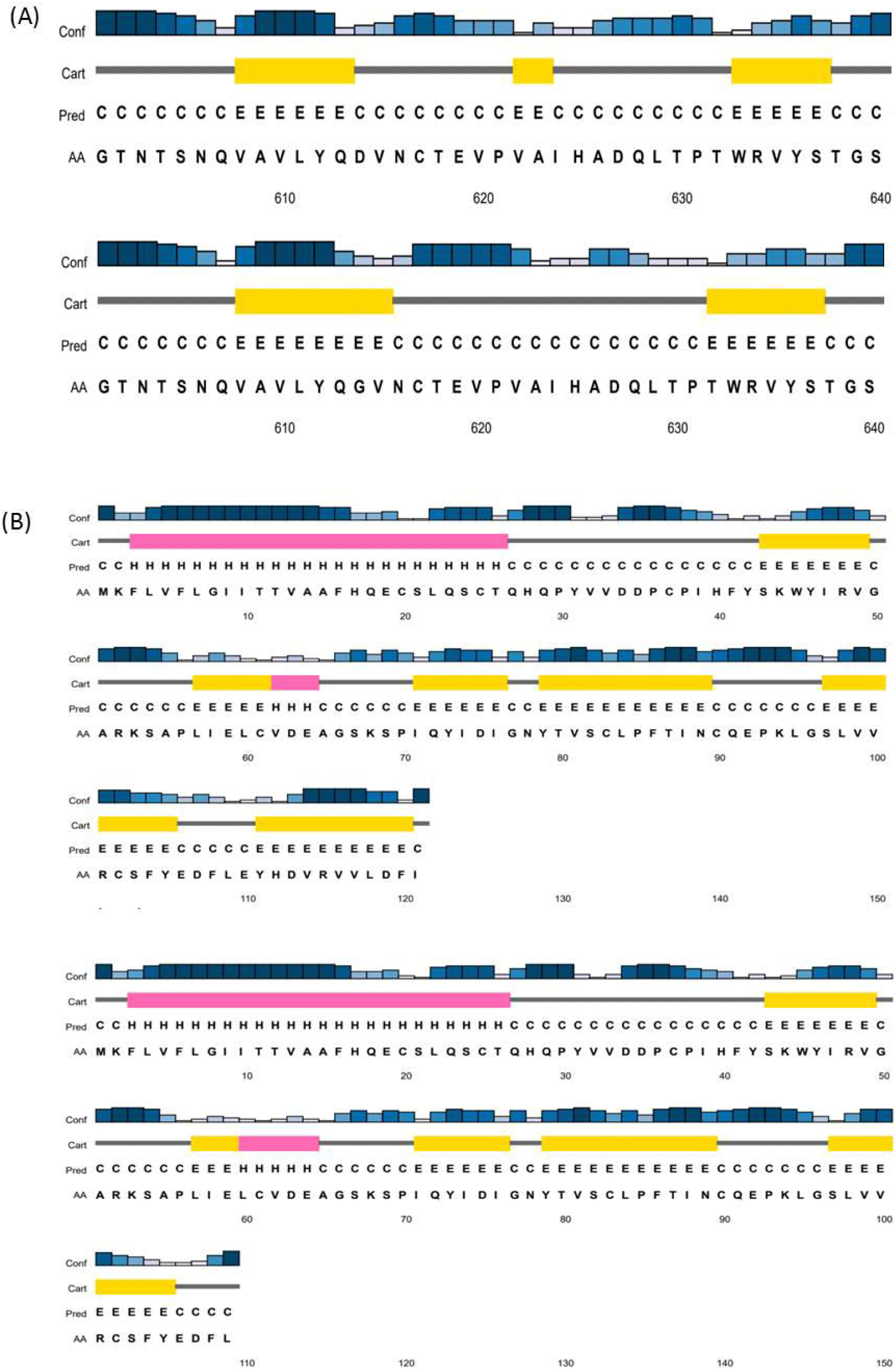
The predicted secondary structures where the yellow bars represent β-strands (E), pink bars represent α-helices (H) and the grey lines represent the coil regions (C). (a) The (AA) represents the respective amino acids and (Conf) bars show the confidence score of secondary structure prediction. Comparison between the predicted secondary structure of wild-type S protein (upper) and D614G mutated S protein (lower) where the green lines represent β-strands and red lines represent α-helices. (b) The mutation D614G disrupts the β-strand following it. Comparison between the predicted secondary structure of wild-type ORF8 protein (upper) and E110stop mutated ORF8 protein (lower). The E110stop mutation essentially truncates off the last beta-strand (110^th^ to 118^th^ amino acid) in the secondary structure of the protein.

As a result of the mutation of G25563T, glutamine gets substituted into histidine in the ORF3a protein. The residues governing this glutamine at position 57 of the ORF3a protein are ^52^L-L-A-V-F-**Q**-S-A-S-K^61^. Histidine substituting glutamine would be converting this sequence site into a more polybasic site that might result in a proteolytic cleavage site. The mutation lies in the loop region of two of the three transmembrane domains of ORF3a viroporin. Since none of the transmembrane domainsis being disturbed, the function of this viroporin might not be affected by this mutation.

Due to the mutation of G28221T, encoding amino acid glutamic acid gets substituted by a stop codon resulting in a 36-nucleotide long deletion in the C-terminal of ORF8. The deletion leads to truncation at the C-terminal of ORF8 which results in a protein of 109 residues instead of 121 of the wild-type protein. Secondary structure prediction of the wild-type protein and the mutated variant (Fig. 2 B) suggests that the C-terminal of this protein (70-118 amino acids) in the wild-type ORF8 protein was found to have four uniformly-spaced beta-strands. However, this mutation was found to essentially chop off the fourth beta-strand. The highly organized secondary structure of the C-terminal of the protein is likely to have some function in the viral replication or infectivity. This mutation truncating the fourth beta-strand probably disturbs its function. Thus, this mutation might be severely affecting the functionality of the ORF8 protein. We also found this same mutation in the virus isolate from an unrelated individual suggesting that this mutation might be relatively common. This result has a striking similarity with that of an in-vivo experiment reported earlier on SARS CoV-1 where a 29-nucleotide deletion in the ORF8 gene was the most obvious genetic change in the early stage of human-to-human transmission of SARS CoV-1[9]. Moreover, the 29-nucleotide deleted SARS CoV-1 strain had a 23-fold less viral replication as compared to its wild type, suggesting that this mutation effectively attenuated the virus. The conspicuous similarity with our result might shed light into the attenuation of the Indian strain of SARS-CoV-2 in humans. It would be interesting to further explore its possibility for vaccine development.

Interestingly, the mutation in ORF8 was found when A at 28254^th^ position was found to be deleted. However, another stop codon was formed 12 nucleotides farther due to this frameshift mutation. As a result, the last two residues of the protein get changed and additional four amino acids get incorporated in the C-terminal of the resulting protein. Since the modifications are at the terminal and in a non-conserved region of the protein, this mutation is unlikely to affect the activity of ORF8.

The mutation of G28371T results in the substitution of serine into isoleucine which lies in a region away from any of the functional domains of the N protein. Thus, it is less likely to have any significant effect on the structure of the protein. However, with the amino acid residues ^18^G-G-P-S-D-S-T-G-**S-**N-Q-N-G-E^31^ around this position 26 of the N protein, mutation of serine into isoleucine is likely to bury the protein structure inside due to the hydrophobic nature of isoleucine. As predicted by NetOGlyc 4.0 Server, we find this region to be a potential O-linked glycosylation site. All the serine and threonine residues mentioned in the short surrounding sequence above are predicted to be a site for O-linked glycosylation making it a heavily glycosylated site. Since glycosylation of viral proteins is a very important post-translational modification governing the virulence of a virus, mutation of this site into a hydrophobic residue can compromise the glycosylation process. Presence of glycine and proline residues around this region is expected to give the structure kink and expose these residues for a potential O-linked glycosylation process. Hence, this mutation of serine into isoleucine is expected to heavily inhibit and compromise this modification in this region.

## Acknowledgement

We would like to thank the Department of Science and Technology, Government of Gujarat for financial support.

